# Effect of Lys84 carbamylation and chloride ion on OXA-143 dynamics and catalytic efficiency

**DOI:** 10.1101/2025.10.27.684880

**Authors:** Víctor U. Antunes, Yara Lins Rocha, Rafael Rospendowski, Luiz De Martino Costa, Renan A. S. Pirolla, Fabio C. Gozzo, Ronaldo J. Oliveira, Denize C. Favaro

## Abstract

Carbapenem-hydrolyzing Class-D β-lactamase OXA-143 is known as an efficient carbapenemase, and the number of clinically isolated *Acinetobacter baumannii* carrying the OXA-143 gene has risen significantly since it was first identified in 2004. The inhibitory effect of chloride ions on OXA enzymes has been described in the literature; however, no study has been performed on the OXA-143 subfamily. Furthermore, there is no information available on the effect of carbamylation on the OXA dynamics. Here, we explored the effects of the absence and presence of carbamylated lysine on protein dynamics using Mass Spectrometry Hydrogen-Deuterium exchange (HDX) and Molecular Dynamics Simulation (MD). We also investigated the effect of chloride ions on the enzyme kinetics and thermal stability. Our results show that in the absence of the carboxy group, the active site of the enzyme and its surroundings are more flexible due to a lack of hydrogen bonds involving the modified lysine. Furthermore, chloride plays no role in the thermal stability or efficiency of the enzyme in the presence of bicarbonate. However, a decreased efficiency proportional to the chloride concentration was observed in a non-supplemented buffer.

## Introduction

Antibiotics have saved millions of lives. However, their misuse has resulted in the emergence of antibiotic-resistant bacterial strains*(1, 2)*. Bacterial resistance is a result of the ability of these organisms to acquire mutations or obtain genes encoding antibiotic-inactivating enzymes from other bacteria, consequently reducing the efficacy of antimicrobials. Resistance to the β-lactam antibiotics in Gram-negative bacteria is mediated mainly by β*-*lactamases, which, according to their primary sequence and mechanism, are grouped into 4 different classes: A, B, C, and D. Classes A, C, and D are serine-β*-*lactamases (SBLs), whereas those of class B are zinc-dependent enzymes*(3-7)*. In terms of the percentage increase in new enzymes and variants, class D, especially the Carbapenem-hydrolyzing Class-D β-lactamases (CHDLs), has seen the most considerable recent growth, being widely distributed among Gram-negative pathogens, such as *Acinetobacter sp(4, 8-10)*.

Class D enzymes, more commonly known as OXAs, exhibit similar folds and share highly conserved sequence motifs such as the STFK (S^81^-T^82^-F^83^-K^84^) motif, which contains the catalytic S81 and the unusual K84, which, after being carbamylated, works as a general base. The hydrophobic nature of the active site gives lysine 84 an unusually low pKa, enabling the reversible carbamylation to happen – an essential step for the OXA’s catalytic efficiency*(5, 11)*. Nonetheless, the great majority are focused on the effect of bicarbonate on protein stability and catalytic efficiency. Another topic frequently discussed in the literature is the inhibitory effect of halide anions on OXA enzymes, with partial inactivation of OXA enzymes reported due to the decarbamylation of the catalytic lysine (KCX)*(12-15)*. Here, we described how carbamylation affects the dynamics and thermal stability of OXA-143 using Mass Spectrometry Hydrogen-Deuterium exchange (HDX), circular dichroism (CD), and MD simulation. We also present Nuclear Magnetic Resonance (NMR) assays that demonstrate the effect of NaCl on the efficiency of the OXA-143 enzyme, present in the absence of bicarbonate.

## Results and Discussion

### Effect of Cl^-^, HCO_3_^-^, and SO_4_^2-^on the enzyme’s thermal stability and secondary structure

To analyze the impact of different bicarbonate concentrations on the thermal stability of the OXA-143, circular dichroism (CD) spectropolarimetry was used to determine the melting temperatures of the enzyme at different anion concentrations (Cl^-^, HCO_3_^-^, and SO_4_^2-^). **Figure 1A** shows that adding bicarbonate increased the thermal stability of the enzyme but not indiscriminately. Inspection of the curves reveals a significant improvement in protein stability when the concentration of NaHCO_3_ increases from 0 to 25 mM and from 25 to 200 mM. Very little gain of stability is obtained at concentrations above 25 mM bicarbonate. To rule out the effect of ionic strength and to investigate the effect of chloride on the enzyme stability, the assay was also performed in the presence of different concentrations of Na_2_SO_4_ and NaCl. **Figure 1B and 1C** show that the protein stability is the same at 0, 25, and 200 mM of sodium sulfate and sodium chloride. These results also eliminate the possibility of any destabilization or decarbamylation caused by the addition of chloride, once the protein is clearly less stable at 0 mM bicarbonate. Also, no changes in secondary structure were observed in the presence of the three ions (**Figure S1**).

**Figure 1.**
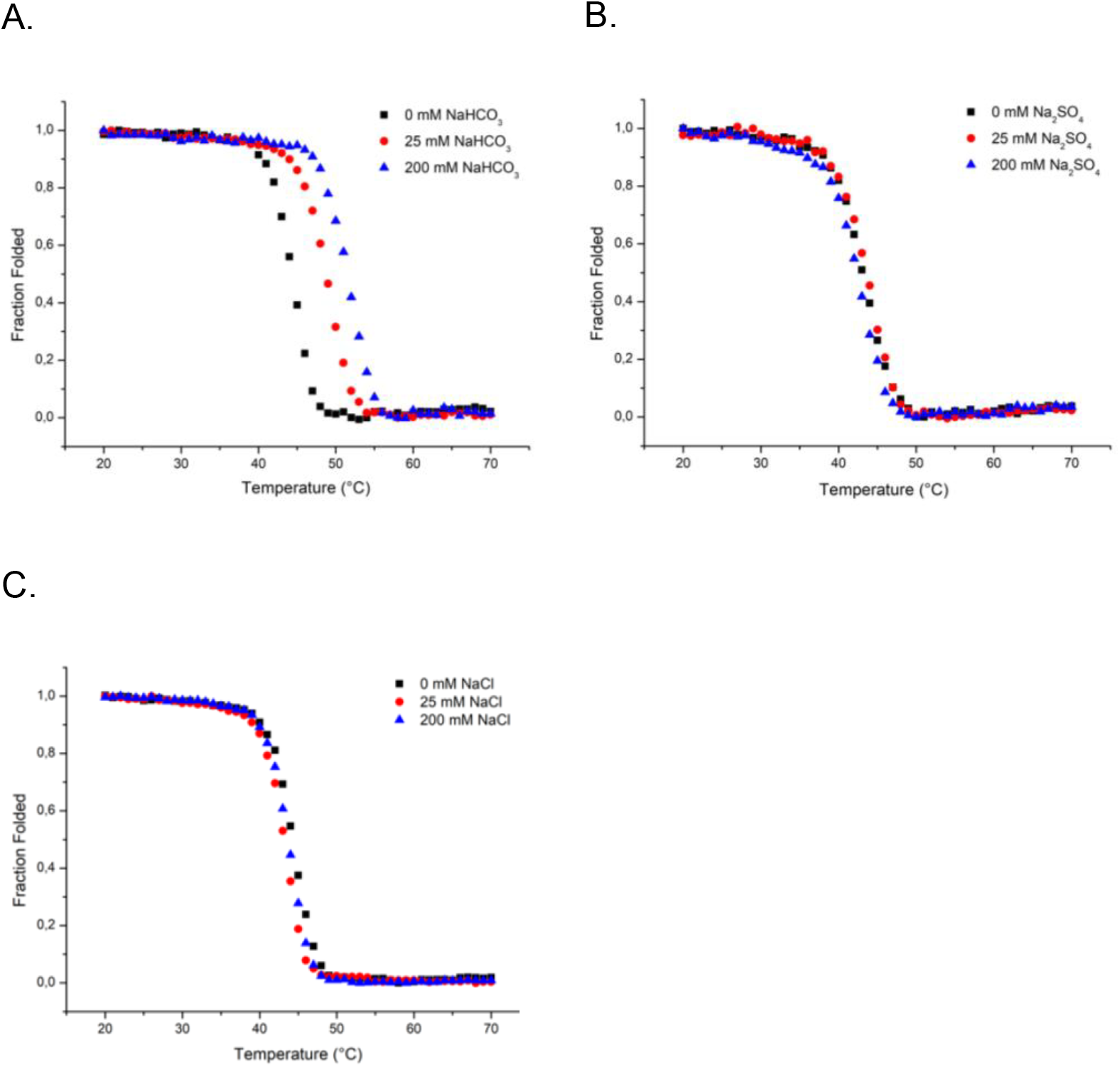
Thermal denaturation (measured at 222 nm) of OXA-143 in the presence of 0, 25, and 200 mM of (A) NaHCO_3_, (B) Na_2_SO_4_, and (C) NaCl.

### Hydrogen-deuterium exchange

The effect of the absence of the carbamoyl group on the protein dynamics was analyzed via hydrogen-deuterium exchange (HDX) mass spectrometry. In **Figure 2** and **Figure S3**, the regions of the protein showing increased exchange in the absence of bicarbonate are highlighted. The peptide LSAVPVYQEL, which includes the conserved S^128^A^129^V^130^ motif involved in product release, followed by the region (H^74^TEYVPASTK^84^), which consists of the active site (S^81^-K^84^), is the ones that show significant differences between the two conditions. The analysis of the interactions performed by the carbamylated lysine, **Figure 2**, shows that the carbamoyl group is involved in hydrogen bonds with S81, W167, and a water molecule. In the absence of this group, these residues would experience a higher mobility, which explains the highest hydrogen-deuterium exchange in the absence of bicarbonate. This may also explain the low thermal stability of the non-carbamylated enzyme.

**Figure 2.**
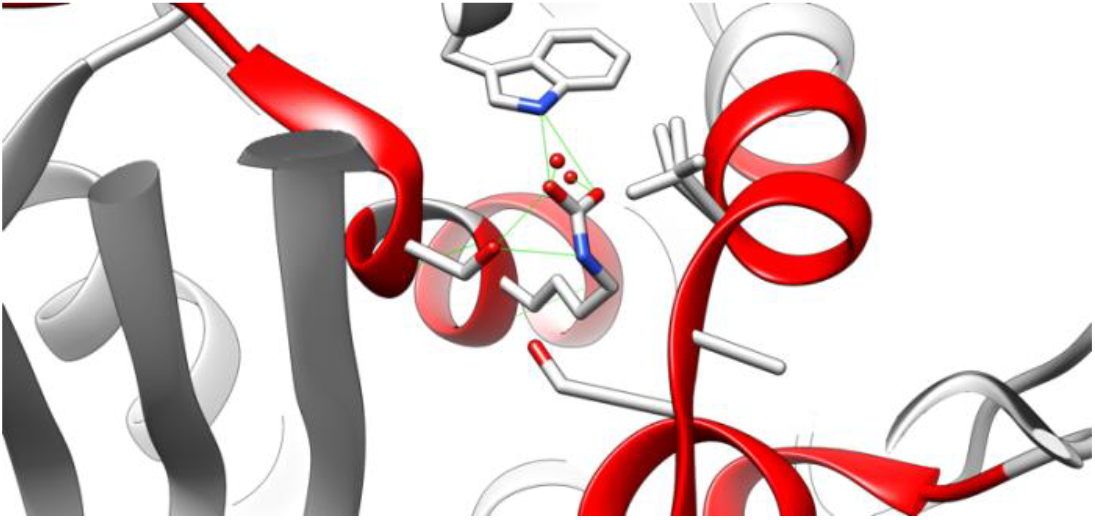
Differences in the hydrogen-deuterium exchange are plotted in the OXA-143 structure. The **red** regions represent where the non-carbamylated OXA-143 presents more deuteration than the carbamylated OXA-143 (PDB: 5IY2).

### MD Simulation

#### Carbamylation enhances the structural and electrostatic stability of OXA-143

All-atom explicit solvent molecular dynamics (MD) simulations were performed on the carbamylated (KCX) and non-carbamylated (K84) forms of the OXA-143 enzyme. The initial structures were energy-minimized and then subjected to 100 ns production runs. Structural stability was evaluated from the fluctuations of two reaction coordinates, the radius of gyration (*R*_*g*_) and root mean square deviation (RMSD), along the trajectories (**Figure S2**). The non-carbamylated enzyme (OXA-143-K84) deviated more strongly from its starting structure, reaching over 4 Å RMSD compared to ∼2.5 Å for the carbamylated form (OXA-143-KCX). By contrast, the *R*_*g*_ values remained similar for both enzymes, with OXA-143-K84 showing slightly higher fluctuations. These results indicate that both enzymes largely preserved their three-dimensional architecture, but the carbamylated form was more structurally stable.

To further probe local flexibility, we computed the root mean square fluctuations (RMSF) of each residue over the MD simulations (**Figure 3**). Consistent with the RMSD analysis, OXA-143-K84 exhibited overall higher flexibility, reflected by increased RMSF values (**Figure 3a**). The P-loop motif, which connects the first half of the helical bundle, was particularly affected by the removal of the carbamoyl group, showing increased fluctuations. This effect is evident both in the primary sequence (**Figure 3b**) and in the projection of RMSF values onto the three-dimensional structure of the enzyme (**Figure 3c**).

**Figure 3.**
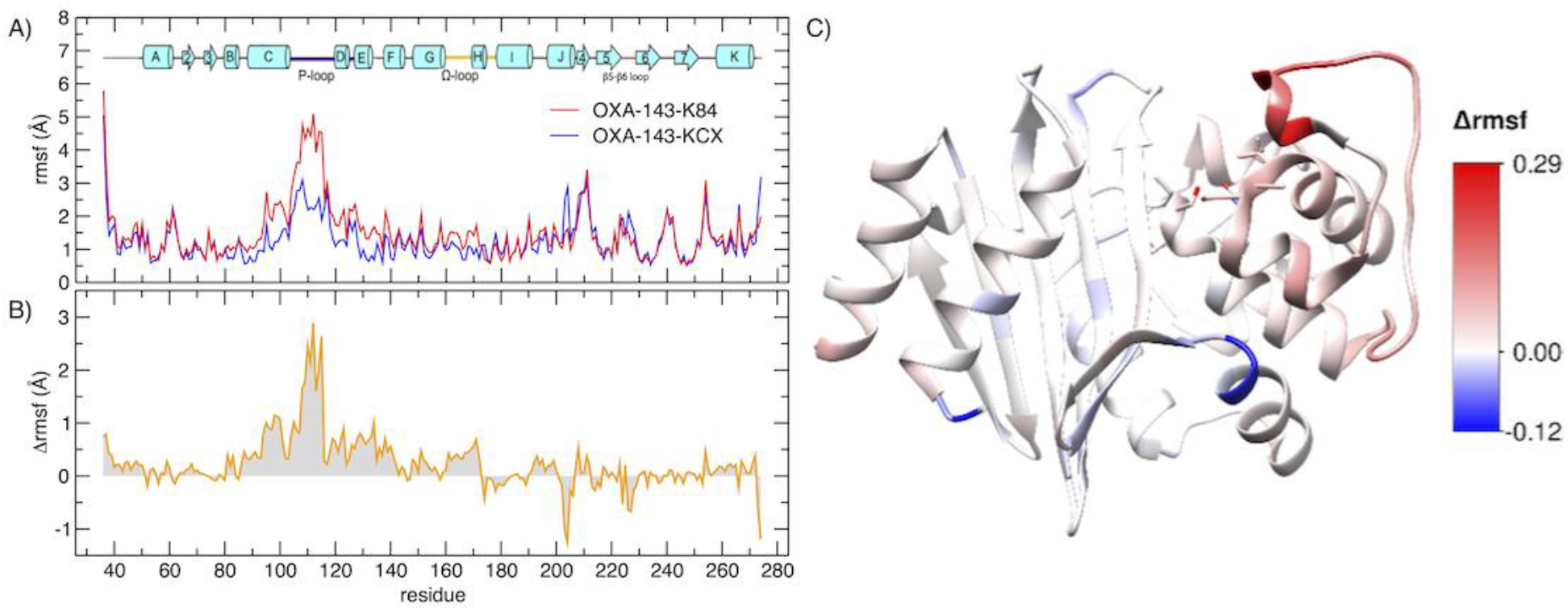
Carbamylation enhances conformational stability and dynamics. **(A)** Root mean square fluctuation (RMSF) per residue for the simulated OXA enzymes. The secondary structure is depicted above the plot. **(B)** Difference in RMSF between the non-carbamylated OXA-143-K84 and the carbamylated OXA-143-KCX. **(C)** Three-dimensional structure of OXA-143 colored by ΔRMSF (RMSF_K84_ − RMSFf_KCX_), with red indicating regions that become more flexible and bluer indicating regions stabilized by decarbamylation. The non-carbamylated enzyme exhibits overall increased flexibility, particularly in the vicinity of the P-loop and its surrounding α-helices.

Complementing the structural and dynamical analyses, the carbamylated (KCX) and non-carbamylated (K84) forms were also examined through electrostatic calculations using the Tanford–Kirkwood solvent accessibility (TKSA) framework*(16, 17)*. TKSA evaluates the contribution of ionizable residues to electrostatic interactions, while accounting for solvent accessibility of each titratable site, and computes the electrostatic free-energy contribution (*ΔG*) of each residue to folding stability. **Figure 4a** shows the electrostatic contribution of each ionizable residue to the stability of KCX and K84 at pH 7. Negative values indicate stabilizing contributions to the native enzyme, whereas positive values indicate destabilizing effects, with red bars highlighting solvent-exposed residues. Removal of carbamylation at position 84 converts KCX back to lysine (K84), directly altering the electrostatic ΔG of neighboring residues, particularly those close to K84. **Figure *4*c** provides a zoomed-in view of the three-dimensional structure surrounding residue 84, where the non-carbamylated form slightly stabilizes the interaction between the solvent-exposed, destabilizing D224 and K84, while eliminating a strong interaction with K218. Overall, carbamylation at position 84 improves the electrostatic contribution to OXA-143 stability, as further supported by **Figure 4b**, where *ΔG* profiles across the pH range reveal stronger stabilizing (more negative) interactions for KCX.

**Figure 4.**
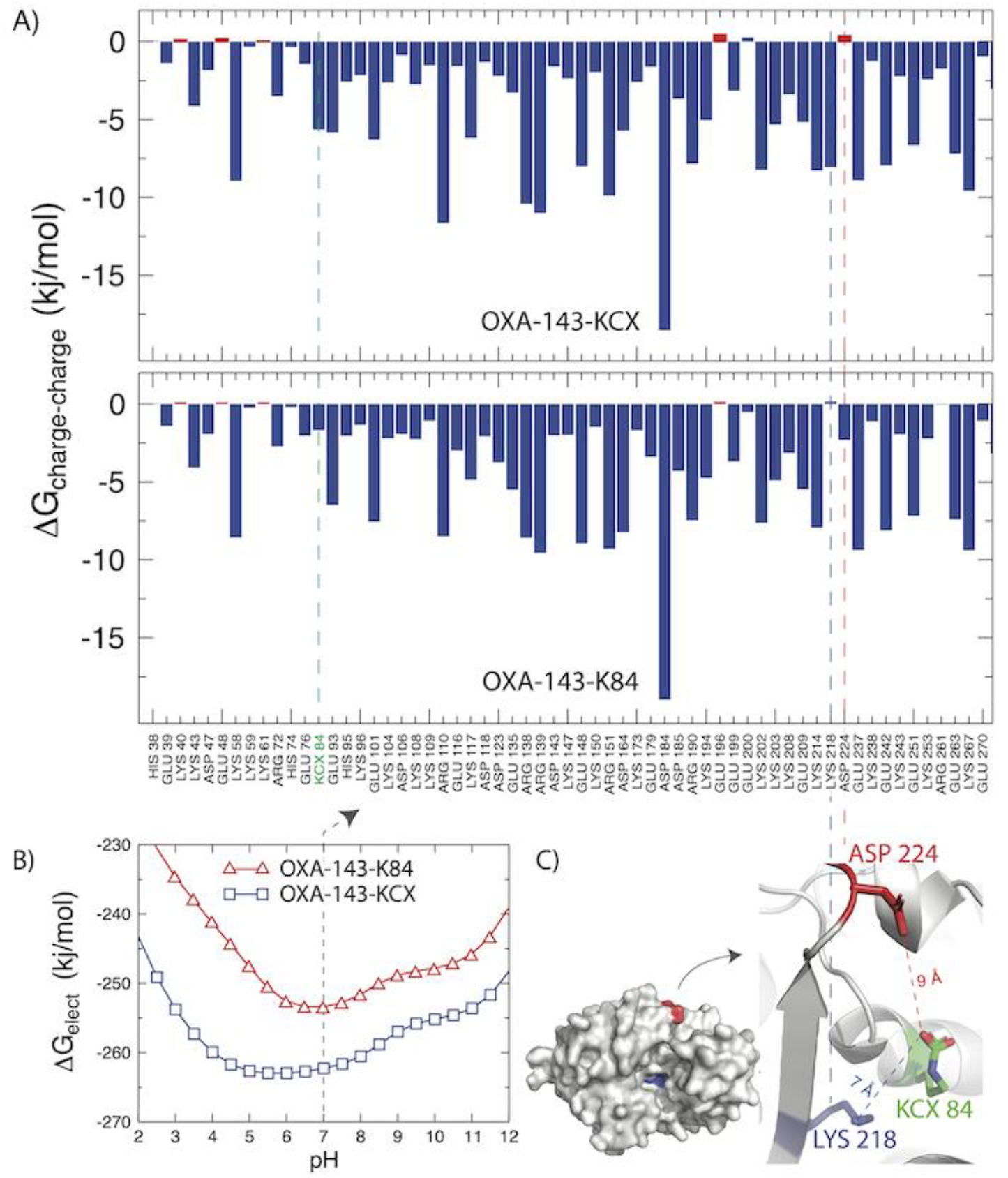
Carbamylation improves electrostatic stability. **(A)** Electrostatic free-energy contribution (ΔG_charge-charge_) of each ionizable residue in OXA-143 variants at pH 7. Bars represent the contribution of each residue to the native-state electrostatic stability relative to the unfolded state. Negative values indicate residues that stabilize the native fold, while positive values correspond to destabilizing contributions. Solvent-exposed destabilizing residues are highlighted in red. **(B)** Total electrostatic free energy (ΔG_elect_) of OXA-143 as a function of pH for the non-carbamylated (K84, red triangles) and carbamylated (KCX, blue squares) forms. **(C)** Three-dimensional structure of OXA-143-KCX highlighting charged residues D224 and K218 in proximity to residue 84. Carbamylation at K84 strengthens electrostatic interactions, particularly with K218, while decarbamylation disrupts this interaction and slightly stabilizes the exposed D224.

### NMR Kinetics

To investigate the effect of chloride on the enzyme’s efficiency, we performed kinetic assays against ampicillin, varying the NaCl concentration from 0 to 300 mM in phosphate buffer supplemented with 25 mM bicarbonate and in a non-supplemented buffer. **Figure 5A** shows that, in the presence of bicarbonate, the efficiency of the enzyme is the same at all chloride concentrations. However, in the absence of supplementation, the reaction slows down in a manner related to NaCl concentration after the first 12 minutes of reaction. Considering that carbon dioxide has low solubility in water, and the enzyme can undergo decarboxylation during the enzymatic reaction, our results suggest that the enzyme is carbamylated at the start of the reaction but is decarboxylated during its course. Furthermore, our UV-Vis assays **(Figure S4)** indicate an increased Km in the presence of NaCl, especially at concentrations of 100 and 300 mM, but no significant effect on the k_cat_. In the presence of a high concentration of chloride ions and the absence of bicarbonate, Cl^-^ may form the ion-pair with residues R261 and S81, similarly to what happens to the OXA-48 *(14)*, decreasing the enzyme-substrate affinity.

**Figure 5.**
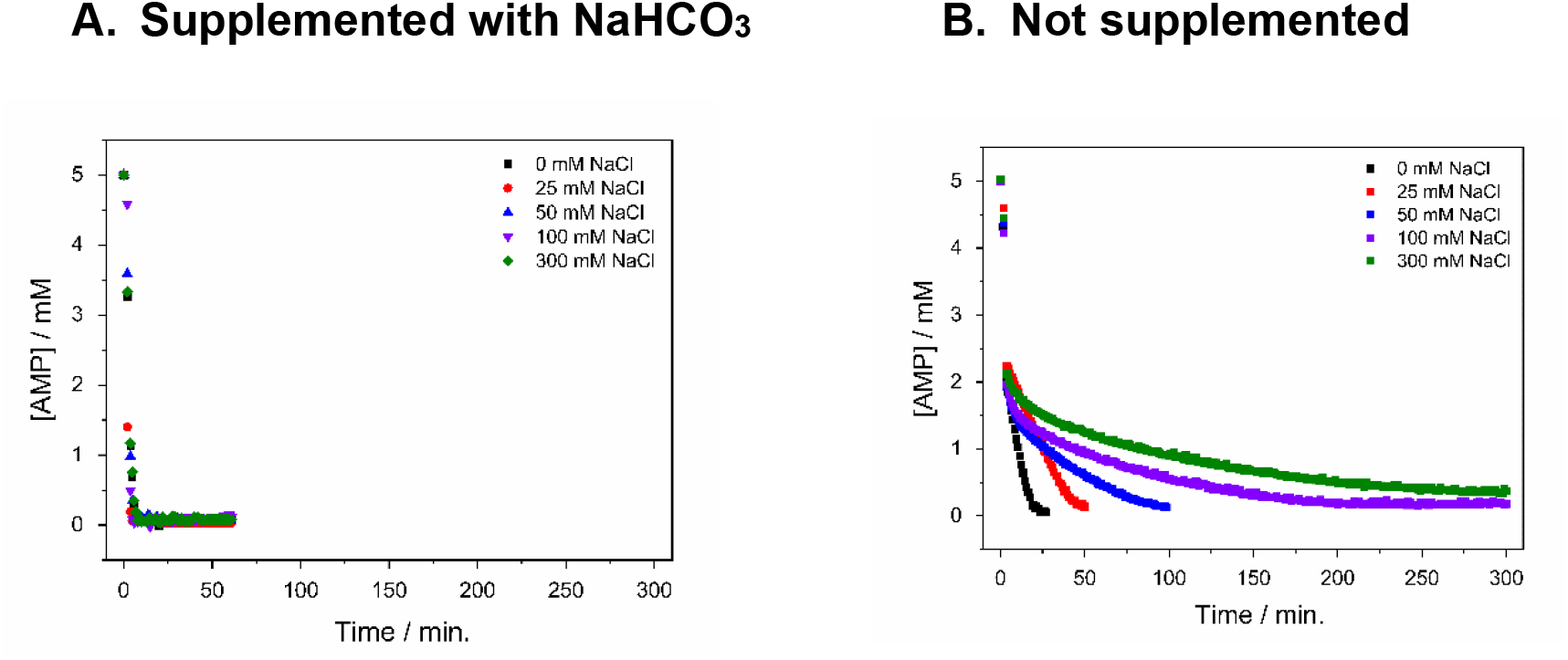
Time-course hydrolysis of 5mM of ampicillin (AMP) in 20 µM OXA-143. Experiments were performed in 50 mM phosphate, 25 mM NaHCO_3_, pH 7.0, with varying NaCl concentrations of 0, 25, 50, 100, and 300 mM. (A) buffer supplemented with 25 mM NaHCO_3_. (B) 0 mM NaHCO_3_.

Furthermore, we also analyzed the effect of NaCl on the stability of the carbamate. **Figure 6A** shows the stacked 13C-NMR spectra of 25 mM NaH^13^CO_3_ (blue), OXA-143 + 25 mM NaH^13^CO_3_ (red), and OXA-143 + 25 mM NaH^13^CO_3_ + 300 mM NaCl (green). No decarbamylation is observed in the presence of high salt concentration, which agrees with our kinetics results, which show that chloride plays no role in the enzyme efficiency in the presence of bicarbonate. Moreover, we tested for the possibility of decarbamylation in the presence of both a high concentration of sodium chloride (300 mM) and 10 mM of the inhibitor tazobactam (**Figure 6B)**. Again, no effect is observed, suggesting that in vivo, where there’s always bicarbonate present, chloride plays no role in the enzyme activity.

**Figure 6.**
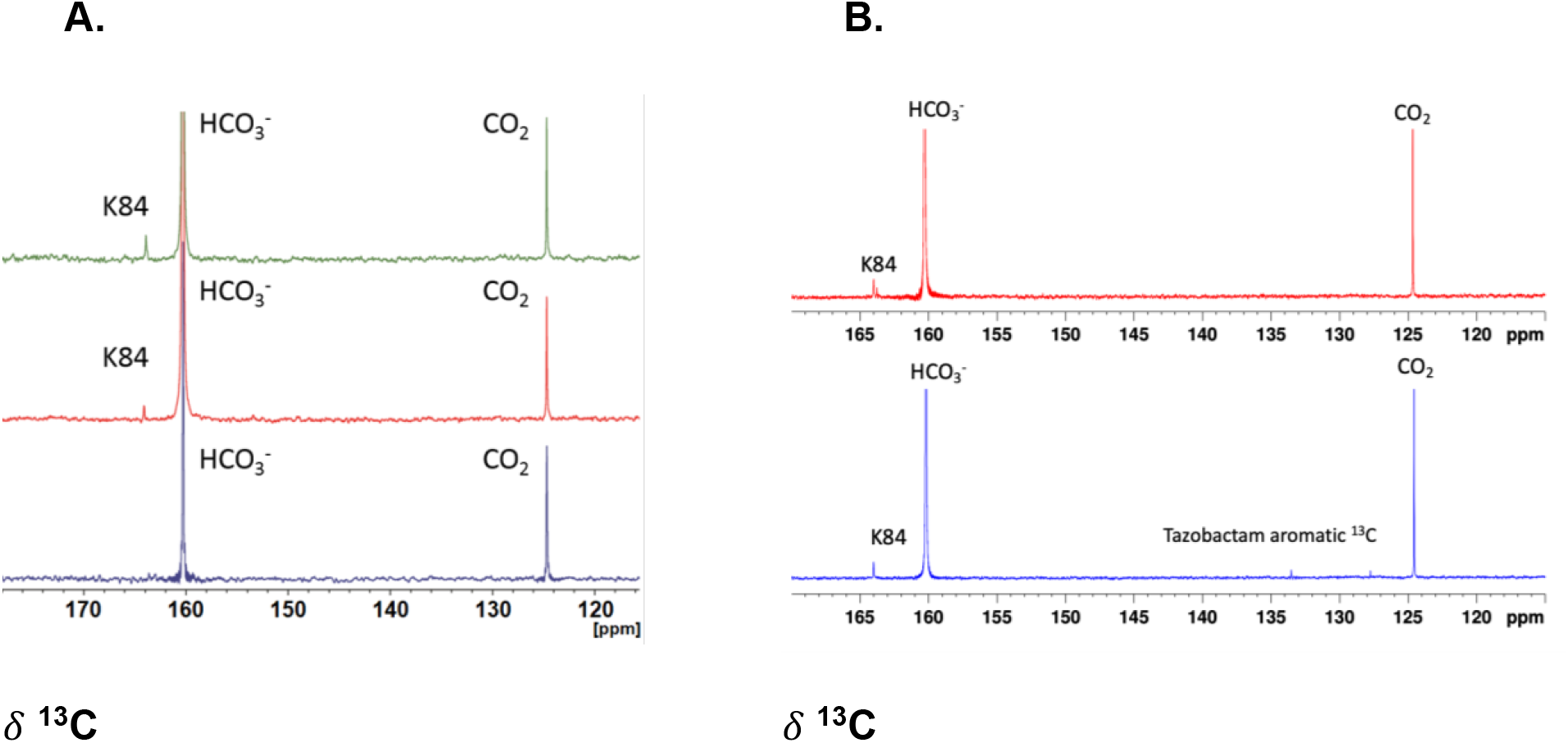
^13^C NMR spectrum **(A)** Blue: 25 mM NaH^13^CO_3_; red: 280 µM OXA-143 + 25 mM NaH^13^CO_3_; green: 280 µM OXA-143 + 25 mM NaH^13^CO_3_ + 300 mM NaCl. **(B)** Blue: 280 µM OXA-143 + 300 mM NaCl + 10 mM Tazobactam + 50 mM NaH^13^CO_3_; red: 280 µM OXA-143 + 300 mM NaCl + 50 mM NaH^13^CO_3_.

## Conclusion

This study examines the role of K84 carbamylation in the structural, electrostatic, and functional stability of the class D β-lactamase OXA-143. Using experimental and computational methods, we demonstrate that bicarbonate (HCO_3_−) improves the enzyme’s thermal and conformational stability, whereas chloride (Cl−) and sulfate 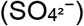 anions do not affect the enzyme’s thermal stability or secondary structure, indicating that chloride does not cause destabilization or decarbamylation.

Hydrogen-deuterium exchange (HDX-MS) revealed that the absence of carbamylated lysine results in increased flexibility of the enzyme’s functional regions, including peptides located in the active site (S^81^-K^84^) and the product release region (S^128^A^129^V^130^). Structural analyses confirmed that the carbamoyl group forms stabilizing hydrogen bonds with residues S81, W167, and a water molecule, the loss of which compromises the proper orientation of the groups during catalysis and the thermal stability of the enzyme.

Molecular dynamics simulations further corroborated these results, demonstrating that the carbamylated form (OXA-143-KCX) exhibits lower structural deviation (RMSD), reduced local flexibility (RMSF), and greater conformational stability compared to the non-carbamylated form (OXA-143-K84). Although both forms maintain a similar fold (R_g_), carbamylation makes the enzyme structure more stable. Electrostatic simulations revealed that carbamylation improves the electrostatic folding energy (*ΔG*), favoring stabilizing interactions and reducing the exposure of ionizable residues to the solvent. The absence of the carbamoyl group disrupts these interactions, weakening the electrostatic environment around K84.

Furthermore, our kinetic assays demonstrated that, in the presence of bicarbonate, OXA-143 maintains its catalytic efficiency in hydrolyzing ampicillin, regardless of chloride levels, indicating that carbamylation remains stable under physiological conditions. On the other hand, without bicarbonate, a gradual decrease in the reaction rate was observed with increased NaCl concentration, suggesting decarbamylation of the carbamylated enzymes and a possible formation of an ion pair. ^13^C NMR experiments with labeled bicarbonate confirmed that no decarbamylation takes place in the presence of 300 mM NaCl or the inhibitor tazobactam, reinforcing the inability of chloride to decarbamylate the enzyme, corroborating our enzymatic essays.

Finally, these findings reinforce the importance of lysine carbamylation for the OXA-143 catalytic efficiency and advance our understanding of the molecular mechanisms governing the activity of OXA-type β-lactamases. This knowledge has direct implications for the development of therapeutic strategies and more effective inhibitors to combat antimicrobial resistance.

## Materials and Methods

### Protein expression and purification

Bacteria were grown to an optical density of 0.8 at 600 nm, and protein expression was induced using 0.5 mM of isopropyl β-D-1-thiogalactopyranoside (IPTG). After 24 h of expression at 16 °C, the cells were centrifuged (4.000 rpm at 4 °C) and suspended in lysis buffer (50 mM NaH_2_PO_4_ and 10% sucrose at pH 7.0). Soluble proteins were loaded onto a HisTrap (GE) column and washed with 10 column volumes of washing buffer containing 50 mM NaH_2_PO_4_ 20 mM imidazole at pH 7.0. Bound proteins were eluted using elution buffer (washing buffer with 500 mM imidazole). The enzyme was then subjected to size-exclusion chromatography using a HiLoad 16/600 Superdex 75pg (GE Healthcare Life Science) and eluted in buffer (50 mM NaH_2_PO_4_, at pH 7.0). The protein concentrations were calculated using the molar extinction coefficient (ε) of 43,430 M^-1^cm^-1^ at 280 nm, with the ε value estimated with ProtParam (https://web.expasy.org/protparam/).

### Thermal Denaturation and Stability

The enzymes were diluted to 11 µM in 50 mM NaH_2_PO_4_, at pH 7.0, with variable concentrations of NaHCO_3_ (0, 25 and 200 mM). The resulting samples were placed in quartz cuvettes with an optical path of 1 mm, and Circular Dichroism (CD) spectra and thermal unfolding experiments were performed on a Jasco J-720 spectropolarimeter. CD spectra were recorded from 260 to 200 nm at 20 °C. Thermal denaturation assays were performed between 20-75 °C, recording the ellipticity at a constant wavelength of 222 nm. All data were analyzed using Origin software.

### Kinetics and analysis of the effect of NaCl on carbamate stability by NMR

Nuclear Magnetic Resonance experiments were performed on Bruker Avance-III spectrometers operating at 600.17 MHz for ^1^H and 125.71 MHz for ^13^C. Antibiotics hydrolysis was monitored at 25 °C using a pseudo-2D pulse sequence with water presaturation during the recycle delay (d1). To analyze the effect of NaCl on carbamate stability, the enzyme OXA was incubated with NaH^13^CO_3_ in the presence and absence of NaCl and Tazobactam and 1D-^13^C NMR experiments were acquired*(18)*.

### Hydrogen-deuterium exchange

The protein dynamics were evaluated by mass spectrometry. Purified samples were prepared in 50 mM NaH_2_PO_4_ buffer, pH 7.0, with and without 25 mM NaHCO_3_ supplementation. Hydrogen-deuterium exchange was initiated by diluting each sample 15-fold (final concentration of D_2_O = 93.8% v/v) with an identical buffer prepared in D_2_O at 25 °C pD 7.0, corrected by the isotopic effect. At each time point (0 s as the control and 10 s, 1 min, 10 min, 30 min, 60 min and 120 min for deuterated experiments), the exchange reaction was quenched by adjusting the pH to 2.4 with an equal volume of quench buffer (800 mM guanidine chloride, 0.8% formic acid, and 20 mM DTT at pH 2.4). Quenched samples were immediately injected into a nanoACQUITY UPLC System with HDX Technology coupled to a Q Exactive Plus high-resolution mass spectrometer (Thermo Scientific). The online digestion was performed on an immobilized pepsin column (2.0 × 30 mm, Applied Biosystems, USA) for 3 min at 15 °C at a flow rate of 40 μL/min. The resulting peptides were trapped and desalted online using an ACQUITY UPLC BEH C18 1.7 μm VanGuard Precolumn (Waters Corp.) at 0 °C. Trapped peptides were eluted into an ACQUITY UPLC BEH C18 1.7 μm, 1 mm × 100 mm column (Waters Co.) equilibrated at 0 °C. Peptides were separated over a 13-minute linear acetonitrile gradient (8–40%) containing 0.1% formic acid at a flow rate of 40 μL/min.

The pH of the aqueous mobile phase was adjusted with formic acid to 2.5 to minimize back-exchange. Mass spectrometric analyses were carried out using a HESI source temperature of 150 °C and spray voltage of 3.8 kV, sheath gas flow rate of 6 (arbitrary units), and auxiliary gas flow rate of 10 (arbitrary units). Peptide identification was performed using a nondeuterated sample, whereas deuterium uptake was calculated from deuterated samples using data from nondeuterated runs (retention time, accurate monoisotopic mass, and isotopic pattern). The data were acquired with a mass resolution of 70 K, and scans were performed in the *m/z* range of 200–2000. A nondeuterated sample was used to obtain the peptide list using PatternLab for Proteomics software in HDX mode*(19)*, with 40 ppm for the precursor mass tolerance and pepsin A with semi-specific digestion and 3 missed cleavages. The peptide list from the nondeuterated samples was used as a reference to calculate the deuterium uptake in the labeled samples using Mass Spec Studio software*(20)*. The deuterium incorporation rate for each peptide was calculated using a retention time window of 0.5 min, and the maximum m/z shift was determined. The uptake differences between protein states were deemed significant at the 95% confidence level. Experiments were performed in triplicate.

### Molecular Dynamics Simulations

#### Electrostatic free-energy and molecular dynamics calculations

Electrostatic interactions among the ionizable residues of the enzyme were evaluated using the Tanford–Kirkwood solvent accessibility (TKSA) framework*(16, 17)* at a temperature of 300 K. In this approach, the protein is represented as a spherical cavity immersed in an electrolyte medium modeled via Debye–Hückel theory. Solvent accessibility corrections were applied to each titratable residue, and the electrostatic free energy, ΔG_*N*_(χ), was computed for each protonation microstate χ in the folded protein. The unfolded reference state was assumed to contribute negligibly to electrostatics. Protonation microstates were sampled using a Metropolis Monte Carlo (MC) scheme as implemented in the TKSA-MC webserver*(21)*.

All-atom molecular dynamics (MD) simulations in explicit solvent were carried out using GROMACS 5.1.4*(22)* with the GROMOS 54a7 force field and a 2 fs integration step.

Initial relaxation of each structure involved sequential energy minimizations in vacuo and then in a solvated environment, using steepest descent followed by conjugate-gradient algorithms. The solvated system was embedded in a cubic box containing SPC216 water molecules, neutralized with counterions, and adjusted to a physiological salt concentration of 0.15 M. Equilibration under constant pressure and temperature conditions was performed for 100 ps. Temperature was regulated at 300 K by velocity rescaling with a coupling time of 0.1 ps, while pressure was maintained using standard barostats. After equilibration, 100 ns production trajectories were generated with snapshots saved every 10 ps, and analyses were performed on the thermally equilibrated segment of each trajectory.

## ACKNOWLEDGMENTS

This work was supported by Fundação de Amparo à Pesquisa do Estado de Minas Gerais (FAPEMIG, Grant No. APQ-03197-24) and Conselho Nacional de Desenvolvimento Científico e Tecnológico (CNPq, Grant No. 307992/2022-5). NCC/GridUNESP and CENAPAD-SP provided computational resources. FAPESP: 2020/12687-4. YLR fellowship (CNPq: 164402/2023-3 and FAPEPS: 2022/03706-0).

