## Supplemental Figures for "Effect of Lys84 carbamylation and chloride ion on OXA-143 dynamics and catalytic efficiency"

Contents:

Figures S1-S4

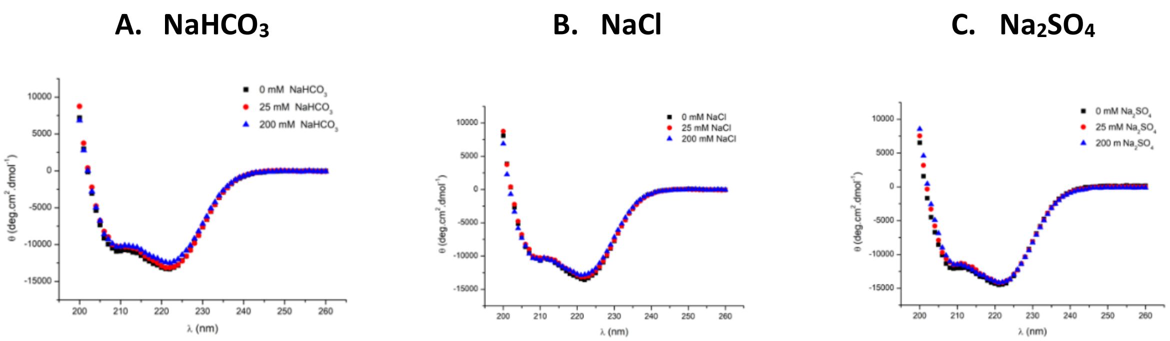


**Figure S1**. Normalized CD spectra of OXA-143 in the absence and presence of 25 and 200 mM of $NaHCO_3$ (A); NaCl (B); $Na_2SO_4$ (C).

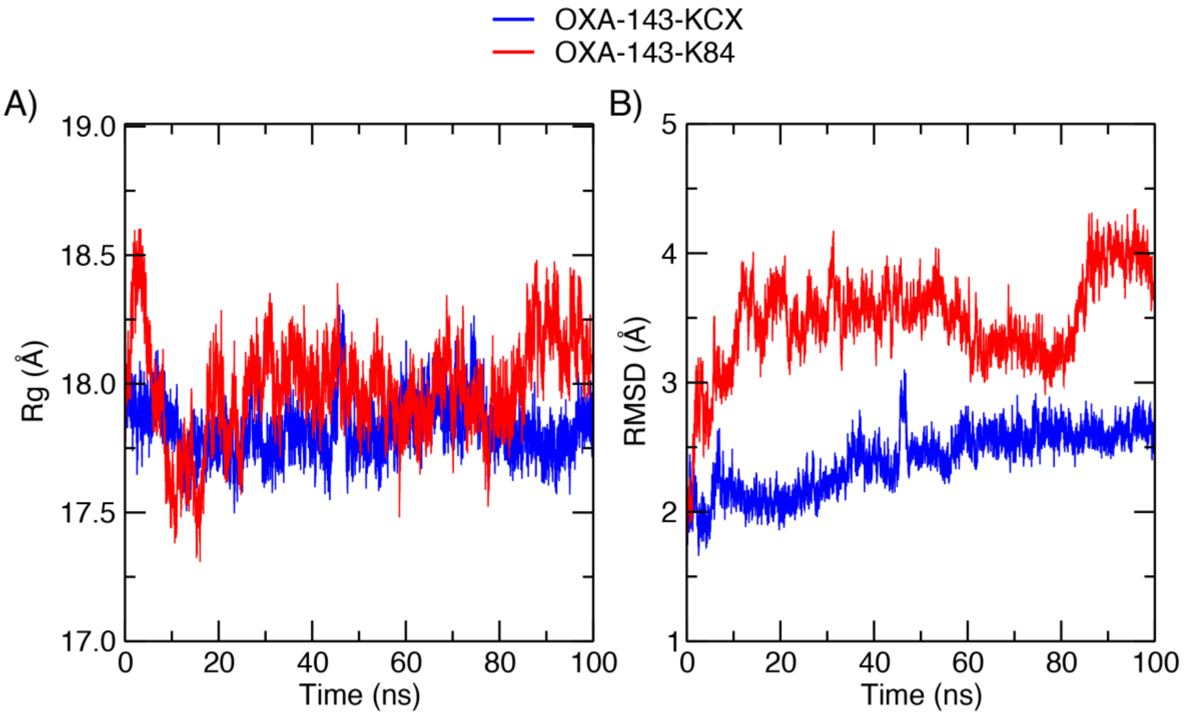


**Figure S2**. Time evolution of (A) radius of gyration ($R_g$) and (B) root-mean-square deviation (RMSD) of the energy-minimized starting structure during all-atom molecular dynamics (MD) simulations. The OXA-143-KCX (blue) maintains a lower $R_g$ and smaller RMSD, indicating a more compact and structurally stable conformation. In contrast, the OXA-143-K84 variant (red) exhibits larger fluctuations and progressive drift, reflecting reduced stability and increased conformational flexibility over the simulated time.

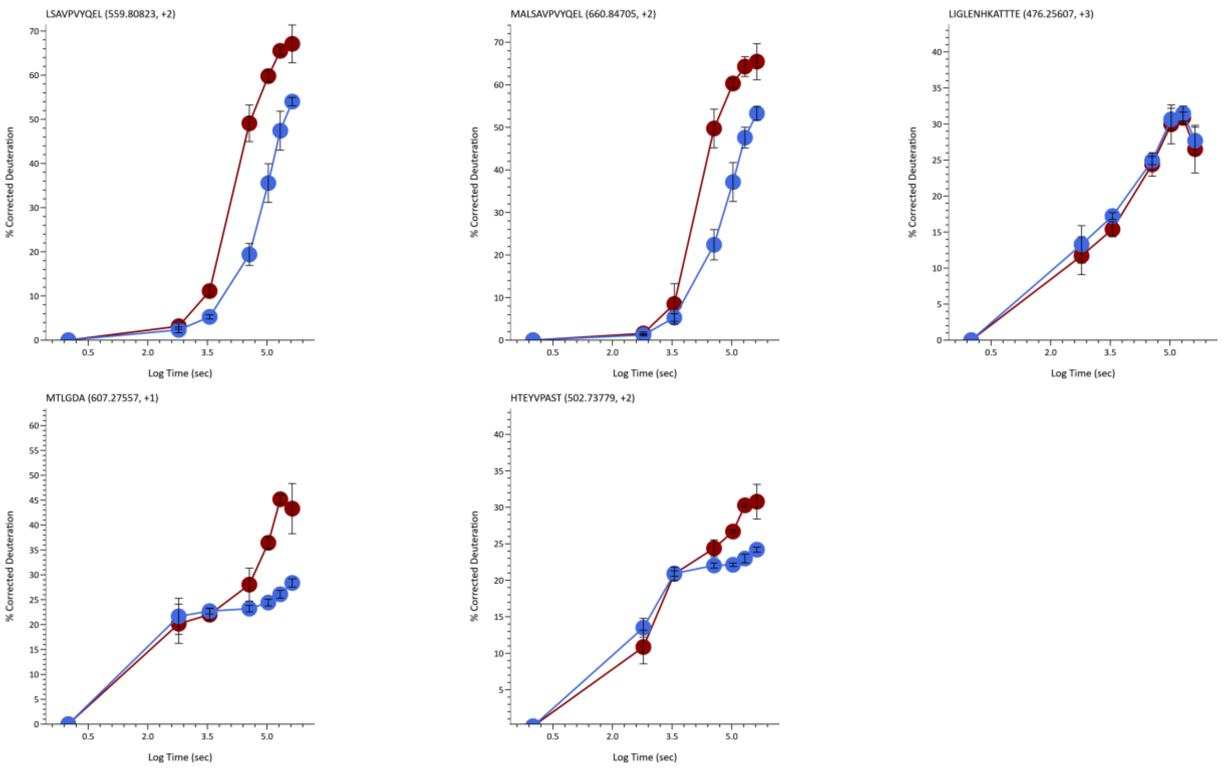


**Figure S3**. HDX peptides: RED – non-carbamylated and BLUE – carbamylated enzyme.

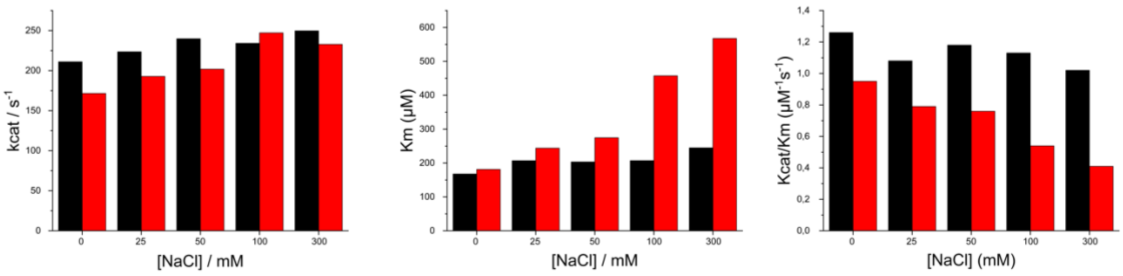


Red: without NaHCO3
Black: supplemented with 25 mM NaHCO3

**Figure S4**. OXA-143 kinetic parameters (kcat, Km, and kcat/Km) against ampicillin at different NaCl concentrations (0, 25, 50, 100, and 300 mM) in the presence (black) and in the absence (red) of 25 mM bicarbonate. The experiments were performed in triplicate, and each value has an average uncertainty of ± 10%.
